# ABRpresto: An algorithm for automatic thresholding of the Auditory Brainstem Response using resampled cross-correlation across subaverages

**DOI:** 10.1101/2024.10.31.621303

**Authors:** Luke A. Shaheen, Brad N. Buran, Kirupa Suthakar, Seth D. Koehler, Yoojin Chung

**Author notes:** Corresponding author: Luke A. Shaheen. Kirupa Suthakar: National Institute on Deafness and Other Communication Disorders, National Institutes of Health, Bethesda MD.

## Abstract

The auditory brainstem response (ABR) is an essential diagnostic indicator of overall cochlear health, used extensively in both basic research and clinical studies. A key quantification of the ABR is threshold, the lowest sound level that elicits a response. Because the morphology of ABR waveforms shift with stimulus level and the overall signal-to-noise ratio is low, threshold estimation is not straightforward. Although several algorithmic approaches have been proposed, the current standard practice remains the visual evaluation of ABR waveforms as a function of stimulus level.

We developed an algorithm based on the cross-correlation of two independent averages of responses to the same stimulus. For each stimulus level, the individual responses to each tone-pip are randomly split into two groups. The median waveform for each group is calculated, and then the normalized cross-correlation between these median waveforms is obtained. This process is repeated 500 times to obtain a resampled cross-correlation distribution. For each frequency, the mean values of these distributions are computed for each level and fit with a sigmoid or a power law function to estimate the threshold.

Algorithmic thresholds demonstrated robust and accurate performance, achieving 92% accuracy within ±10 dB of human-rated thresholds on a large pool of mouse data. This performance was better than that of several published algorithms on the same dataset. This algorithm has now fully replaced the manual estimation of ABR thresholds for our preclinical studies, thereby saving significant time and enhancing objectivity in the process.

## 1. Introduction

Auditory Brainstem Response (ABR) thresholds serve as a translational estimate of hearing sensitivity, providing a crucial metric in basic science, preclinical research, and clinical audiology. Traditionally, the standard in the field has been for expert human raters to use their knowledge and experience to visually determine these thresholds. Over time, there have been numerous attempts to automate this process by developing algorithms to decide the threshold, but none of these attempts have been widely adopted. Some of these algorithmic approaches include assessing waveform similarity to templates (Cone-Wesson et al., 1997; Elberling, 1979), evaluating waveform stability across repeated measures or levels (Berninger et al., 2014; Ozdamar et al., 1994; Suthakar and Liberman, 2019; Wang et al., 2021; Xu et al., 1995), analyzing signal quality through F-ratios and other metrics (Cebulla et al., 2000; Don and Elberling, 1994; Sininger, 1993) and using Bayesian nonlinear regression (Gaussian process, Chesnaye et al. 2024). Other methods have involved neurophysiological parameters derived from fitting the responses to different stimulus intensities (Schilling et al., 2019). More recently, artificial intelligence, in particular deep learning techniques, have been explored as a potential solution, given their ability to infer complex patterns in large datasets (Acir et al., 2006; Erra et al., 2024; Thalmeier et al., 2022). Despite these efforts, the field continues to rely heavily on human expertise for ABR threshold determination.

The automation of ABR threshold estimation is complicated by several factors. Firstly, ABR morphology, or waveform shape, is not static but varies with the frequency and level of the sound stimulus. It also varies based on other factors such as species, strain, genetic model, electrode position, repetition rate, and more. This variability is a challenge to developing a universally applicable algorithm, particularly those that measure waveform similarity across levels or to a template. Secondly, ABRs typically have a low Signal-to-Noise Ratio (SNR), making the estimation of the true threshold difficult. If the ABR has more noise than typical the SNR is decreased, potentially causing an elevated estimation of threshold by a human rater. Because of the low SNR and variable morphology, an iso-response strategy like that used for detecting the thresholds of distortion product oto-acoustic emissions (DPOAE), where the threshold is defined as the stimulus level at which the signal crosses a criterion, is difficult to implement. The ABRpresto algorithm described here transforms the waveform into a single measure for each stimulus level and frequency, allowing for an iso-response strategy. Lastly, even when peak amplitudes are within the noise floor, consistent responses can still be apparent. This means that even low-amplitude signals within the background noise fluctuations, which might be considered noise in other contexts, can be genuine auditory responses. On the other hand, there are also cases where noise peaks that occur in line with ABR peaks at higher stimulus levels are not genuine auditory responses, but are mistaken as such by the human rater, causing underestimation of the threshold.

We developed an algorithm (ABRpresto) for the determination of thresholds based on the resampled cross-correlation of two independent averages of responses to the same stimulus. The algorithm mitigates the uncertainty of low-amplitude signals by measuring a distribution of correlations via resampling of the single-trial data. The results were compared to thresholds determined by expert human raters. Algorithmic thresholds were consistent with human-rated thresholds on a large pool of mouse data.

## 2. Materials and Methods

### 2.1. Animals and ABR acquisition

All experiments were approved by the Animal Care and Use Committee of Decibel Therapeutics. A total of 7,857 ABR waveform stacks from 351 mice were analyzed with ABRpresto and compared to human-rated manual thresholds. This dataset includes mice with normal hearing and hearing loss from multiple strains and genotypes, totaling 123,140 unique ABR waveforms (full dataset is available at https://zenodo.org/records/13987792, Shaheen, 2024a). In addition, analogous performance metrics were calculated on Suthakar and Liberman (2019) dataset to facilitate direct comparison with the current algorithm.

ABRs to acoustic tone-pips were recorded under anesthesia (Ketamine 100 mg/kg, Xylazine 10 mg/kg, i.p.). Mice were placed in an acoustic chamber, and body temperature was maintained at 37°C using a feedback-controlled heating system (FHC). Acoustic stimuli were generated at a nominal sampling rate of 100 kHz and delivered through a custom amplifier and dual speaker system based on designs published by the Eaton-Peabody Labs at the Massachusetts Eye and Ear Infirmary (Hancock et al., 2015).

ABRs were recorded with 3 needle electrodes (LifeSync Neuro) inserted into the skin in the dermal layers: (1) a ground electrode near the base of the tail, (2) a recording electrode at vertex along the midline of the skull between the ears, and (3) a reference electrode through the bare skin ventral to the pinna.

Data were collected using a customized version of Psiexperiment, a plugin-based framework for auditory experiments (Buran and David, 2020). Five-millisecond tone-pips with a 0.5 ms rise-fall time delivered at 81/s in alternating polarity were used for frequency-specific measurements of hearing function. Sound levels were tested in 5 dB steps, at either a 15–105 dB, 15–85dB, or 65–105 dB sound pressure level (SPL) range. Stimulus frequencies ranged from 4 to 45.3 kHz in half-octave steps. Stimuli were presented in interleaved order such that a train of tone-pips containing a single presentation of each level and frequency was repeated in an interleaved ramp paradigm (as described in Buran et al. 2020). Electrical signals were recorded through a digitizing amplifier (TDT RA4PA or Medusa4Z). Responses from 512 repetitions were recorded, and individual responses to each trial and averaged responses were stored for further analysis.

### 2.2. Human-rated manual thresholds

To estimate ABR thresholds, the signal was filtered from 300 to 3000 Hz, and the lowest sound level that elicits an ABR was estimated using custom peak-picking software (EPPeak). Analysis was performed by blinded raters. Most waveform stacks were evaluated by a single rater, but in cases where multiple raters evaluated a waveform stack, the median value was taken as the estimated threshold.

### 2.3. ABRpresto: Automatic thresholding algorithm

Figure 1 is a flow chart of the automatic thresholding process based on cross-correlation distribution to provide more accurate and reliable results. Due to how the processing workflow was configured, single-trial data were filtered twice, forward and backward each time using a first-order 300–3000 Hz Butterworth filter and the *filtfilt* function of Python’s SciPy module (for a total of 4 times). For each iteration: 1) ABR waveform samples are randomly split into two groups, each containing an equal number of responses to positive and negative polarity stimuli; 2) a median waveform for each group is calculated; 3) a normalized cross-correlation between the two median waveforms at a lag of 0 ms is performed to determine a Pearson correlation coefficient. This process is repeated 500 times to obtain a distribution of correlation coefficients for each sound level.

**Figure 1.**
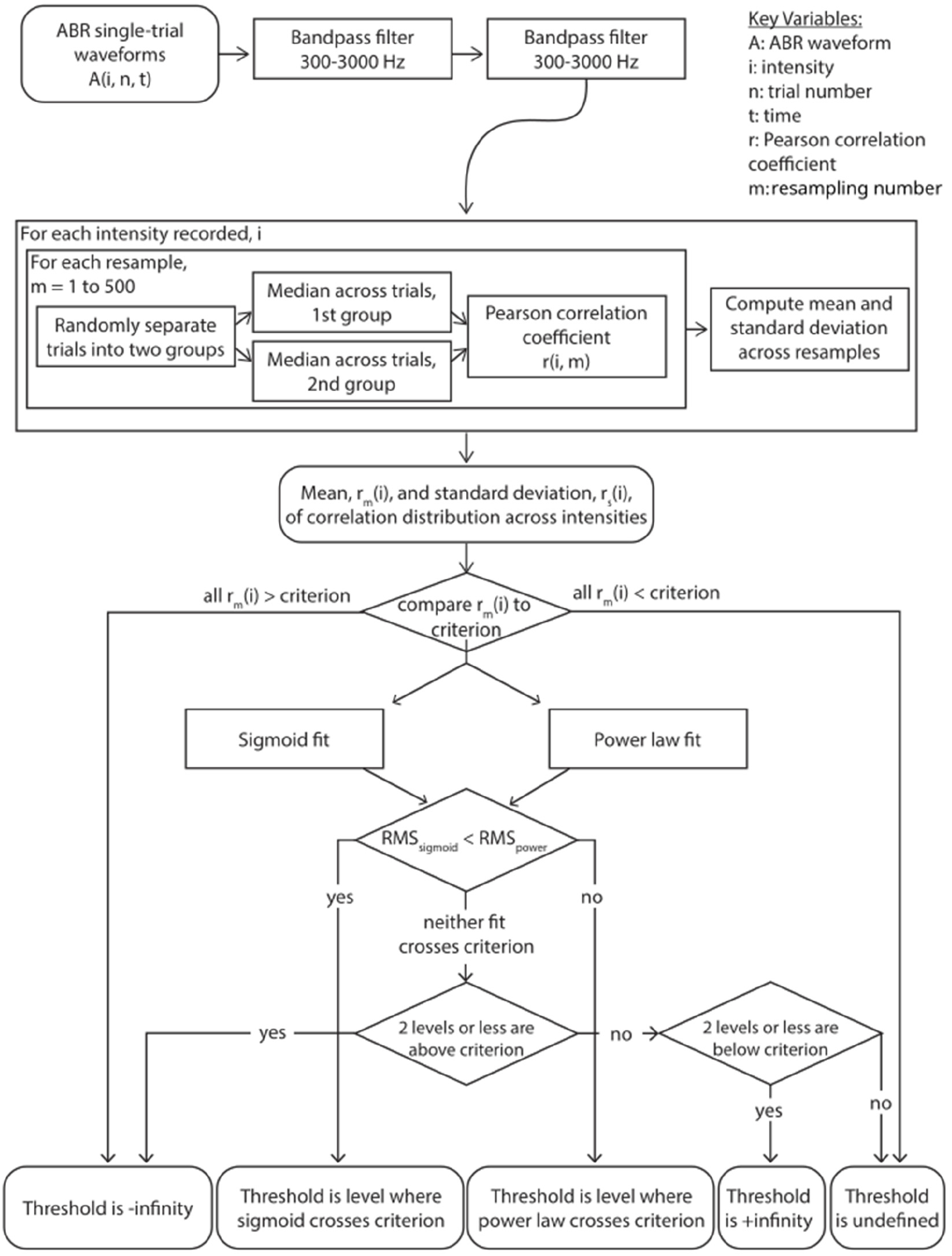
Flowchart of the ABRpresto automatic thresholding algorithm

The mean of the cross-correlation function is then fit with two separate functions: a sigmoid and a power law function, and the best fit is then chosen based on root-mean-squared (RMS) error in a similar way to Suthakar and Liberman (2019). If all means are greater than the criterion, the fits are not considered and the threshold is set to negative infinity as an indicator that the exact threshold is not known, but is below the minimum level tested for that waveform stack. If all means are less than the criterion, the fits are not considered and the threshold is set to positive infinity as an indicator that the exact threshold is above the maximum level tested. If the RMS error of the sigmoid fit is lower, threshold is defined as the level where the sigmoid curve crosses the criterion; otherwise, it is defined by the crossing of the power law curve.

In the event that neither fit crosses criterion, the threshold is marked as undefined for review by a human rater with two exceptions. First, if the means for two levels or fewer are above the criterion, assume that spurious correlations pushed these levels above the criterion and infer that the true result is likely all less than the criterion, but some noise pushed some above. Therefore, we set the threshold to negative infinity. Second, if the means for two levels or fewer are below the criterion, then we set the threshold to positive infinity by the same logic. If neither condition is met, there was an error in fitting and the threshold is set to undefined for further review.

We explored the performance of the algorithm as a function of this criterion (see **3. Results**), and chose a criterion of 0.3. We also explored algorithm performance with and without artifact rejection by excluding trials with any peaks above an absolute level of 20 μV, which excluded trials containing heartbeat artifacts and some larger muscle artifacts. The algorithm performed similarly with and without artifact rejection prior to step 1; the results reported here were computed without artifact rejection. In addition to the mean Pearson correlation coefficient, we evaluated two other metrics to obtain thresholds from the cross-correlation distributions generated by resampled subaverages. 1) Similar to Wang et al. (2021), we measured the cross correlogram as a function of lag between the two subaverages and quantified the percentage of times the peak of the cross-correlation function fell within 0.5 ms. We evaluated several criterion values for this parameter, but none performed well. Unlike Wang et al. (2021), we then fit this percentage with sigmoid and power law functions and found the threshold in the same manner as described above. 2) We quantified the separation between the distribution of cross correlation values at a lag of 0 ms (over resamples) for each level and a null distribution drawn from the lowest tested level using the Kolmogorov-Smirnov test statistic. This value was then plotted as a function of level, fit with a sigmoid and a power law function, and threshold was found as described above. Both approaches yielded worse performance than the ABRpresto method (based on the mean of the cross correlation at a lag of 0 ms).

Figure 2A shows two median ABR waveforms in response to 65 dB SPL tone-pips, and Figure 2B shows the distribution of the cross-correlograms over 500 repetitions (mean ±SD) along with the cross-correlogram of the example shown in Figure 2A. At a sound intensity well above the threshold, the similarity between the two waveforms is apparent, captured by the peak at a lag of 0 ms in the cross-correlogram (Figure 2B). In contrast, two median ABR waveforms in response to 30 dB SPL sub-threshold stimulus do not overlap (Figure 2C). The cross-correlogram distribution at this level shows larger variability and no clear peak at a lag of 0 ms (Figure 2D).

**Figure 2.**
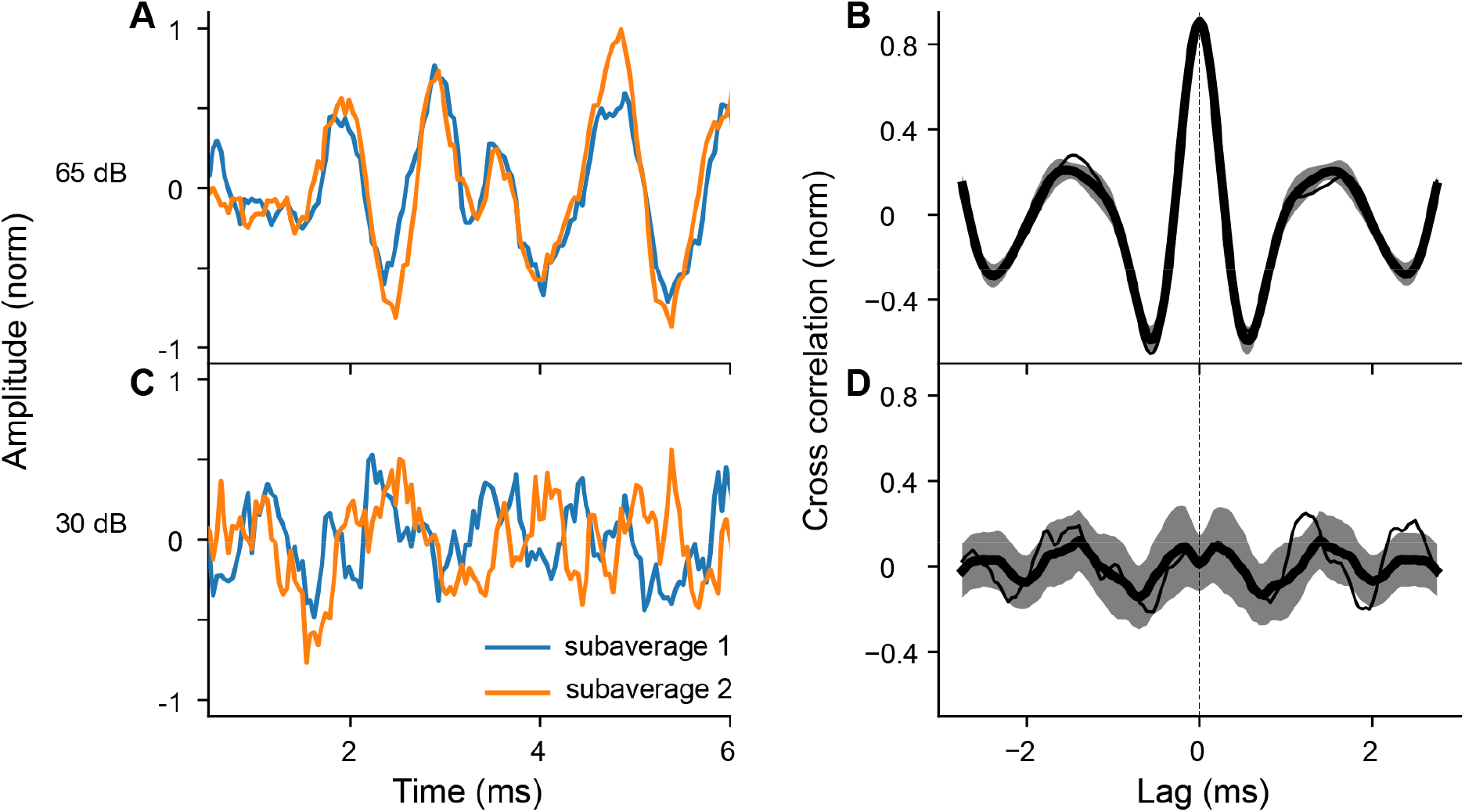
Examples of two subaverages and corresponding cross-correlograms to 8 kHz tone-pips at supra- and sub-threshold sound levels. **A**: Two median ABR waveforms at 65 dB SPL. **B**: Cross-correlograms for all subaverage pairs at 65 dB SPL. Data represented as mean ±SD **C**: Two median ABR waveforms at 30 dB SPL. **D**: Cross-correlograms for all subaverage pairs at 30 dB SPL. For panels B and D, the thin line corresponds to the cross-correlogram of the example pairs in panels A and C, respectively.

Figure 3A shows an ABR waveform stack in response to 8 kHz tone-pips at 15–65 dB SPL. For each sound level, the mean (±SE) of all trials and an example set of two medians from one resample repetition are shown. With increasing sound level, the morphology of ABR peaks becomes clearer, with increasing similarity between the two median waveforms and decreasing overall variability. This trend is clearly captured in the increasing mean and decreasing SD of correlation coefficients shown in Figure 3B. In this example, a sigmoid function results in a better fit, and the threshold is defined as where the curve crosses the criterion of 0.3.

**Figure 3.**
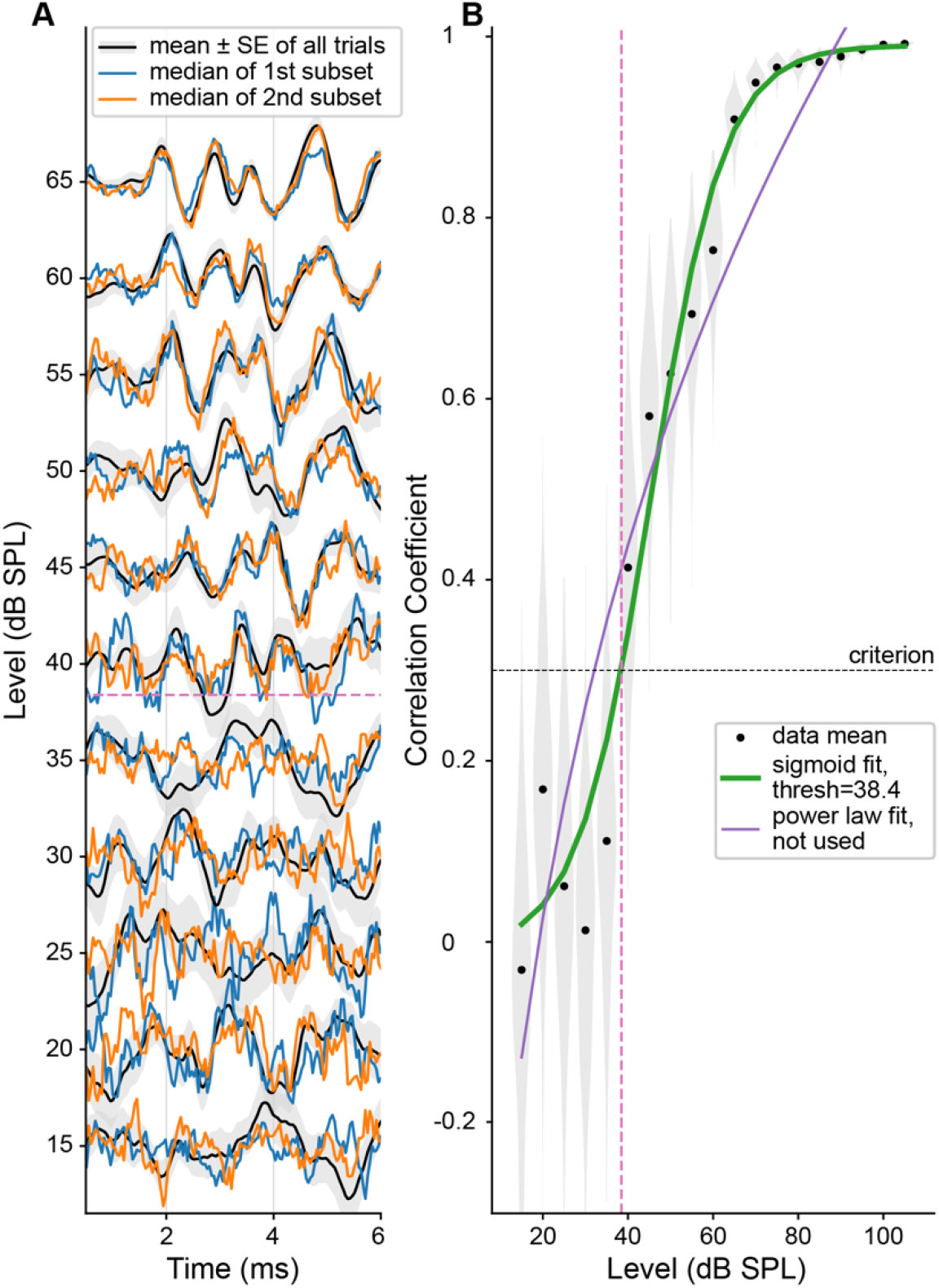
An example fit of ABR presto to 8 kHz tone-pips (same results presented in Figure 2). **A:** Stacked ABR waveforms in response to stimulus levels surrounding the threshold. For each level, mean ±SE from all trials is plotted in black with grey shading, and the medians of the two subsets taken from the first resampling are shown in orange and blue. Waveforms are normalized (for each level, all 3 lines are scaled by the peak-to-peak of the mean of all trials). **B:** Black dots and shading show the mean correlation coefficient and distribution for each stimulus level. Sigmoid and power law fits are shown in green and purple, with the thicker green line indicating that the sigmoid fit performed better, and therefore was used to define threshold. The threshold is shown by the pink dashed line and is defined by where the curve crossed criterion (black dashed line).

### 2.4. Comparison

The algorithmic thresholds were compared to human-rated thresholds. In addition to Spearman’s ρ, the fraction of threshold values within ±10 dB of manual thresholds was computed to capture the absolute difference between human-rated thresholds and algorithmic thresholds. For both methods, when no response was detected within the range of levels tested, the threshold was imputed to a level 5 dB greater than the highest level tested (typically set to 110 dB) and included in the comparison. When a response was detected at the lowest level tested, the threshold was imputed with a level 5 dB lower than the lowest level tested (typically set to 10 dB SPL) and included in the comparison.

The algorithm by Suthakar and Liberman (2019) employs cross-correlation of the average waveforms across levels. We tested this algorithm on the present dataset. Because there was no artifact rejection employed during the collection of the present dataset, but there was in the Suthakar datasets, we applied an artifact rejection prior to averaging across trials by excluding trials with any peaks above an absolute level of 20 μV. We also evaluated the performance of this algorithm as a function of criterion level. Suthakar and Liberman (2019) found that a criterion of 0.35 provided the best performance of their data. On our data, a criterion of 0.6 gave the best performance. The algorithm by Wang et al. (2021) uses cross-correlation across subaverages, but rather than basing the threshold on the value of the cross-correlation at a lag of 0 ms, it instead defines the threshold as the lowest level for which the cross-correlation functions have a peak within ∼1 ms of 0 lag. We re-implemented this algorithm and tested it on the present dataset, again applying 20 μV artifact rejection before averaging.

### 2.5. Code and Data Availability

The code for the ABRpresto algorithm is available for non-commercial use at https://github.com/Regeneron-RGM/ABRpresto (Shaheen, 2024b). The single-trial ABR waveforms used to test the algorithm are available for non-commercial use at https://zenodo.org/records/13987792 (Shaheen, 2024a).

## 3. Results

We created an algorithm to find threshold based on the cross correlation between resampled subaverage (see **2. Material and Methods**). A comparison between manual thresholds vs algorithmic thresholds from 7,857 ABR waveform stacks show a strong correlation (Spearman’s ρ=0.97) between the thresholds estimated manually (by human rater) and by ABRpresto (Figure 4A). Notably, the data set includes a wide range of ABR thresholds from ∼20 dB SPL to no response up to the equipment limit recorded from mice across multiple strains and genotypes and a wide range of frequencies (4–45 kHz).

**Figure 4.**
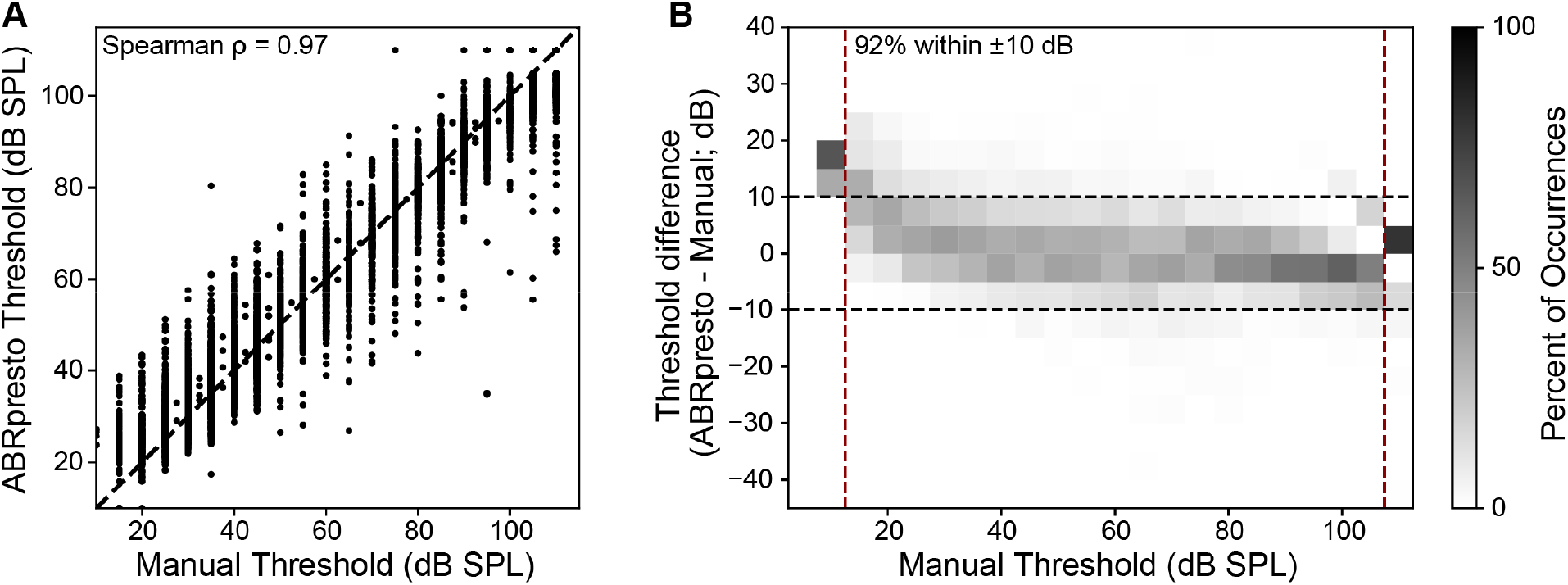
Comparison between manual thresholds and algorithmic thresholds. **A**: Manual threshold vs. algorithmic threshold (Spearman’s ρ=0.97). Each dot represents a single threshold value. **B**: Distribution of threshold difference as a function of manual threshold. Each cell represents the percent of occurrence corresponding to the manual threshold & threshold difference pair, normalized within a column (manual threshold bin). This normalization allows visualization of the distribution of threshold differences as a function of manual threshold. A small bias indicating higher algorithmic thresholds for manual thresholds < 30 dB SPL is apparent. N=7,857 threshold judgements are represented in the figure.

Next, the difference between algorithmic and manual thresholds were computed (Figure 4B). 92% of the algorithmic threshold values fell within ±10 dB of the manual threshold values, indicating strong agreement between the proposed algorithm and human raters. Despite this strong agreement, the algorithm tended to predict higher thresholds than the human rater when the manual threshold was less than 30 dB SPL. In contrast, the algorithm tended to predict slightly lower thresholds when the manual threshold was 30 dB SPL or greater.

Defining the “ground truth” of ABR threshold is a challenging task due to the inherent noise in the evoked potential signal. Figure 5 illustrates an example case for where the human rater likely underestimated the threshold due to noise. The averaged ABR waveform shows clear ABR peaks from 70 dB SPL down to 30 dB SPL. The falling slope of wave 1 is visible in the response to 25 dB SPL. Peaks are observed around the latency matching wave 1 in 20- and 15-dB SPL responses. In this case, the human rater selected 15 dB SPL as the threshold. However, the distribution of median ABR waveforms obtained by resampling shows that peaks in the 15-dB SPL response are not from a robust response and are more likely due to fluctuations in the waveform caused by noise. The low correlation coefficient values over 15–25 dB SPL support this interpretation. The algorithmic threshold in this example is 29.5 dB SPL, a value more than 10 dB higher than the manual threshold.

**Figure 5.**
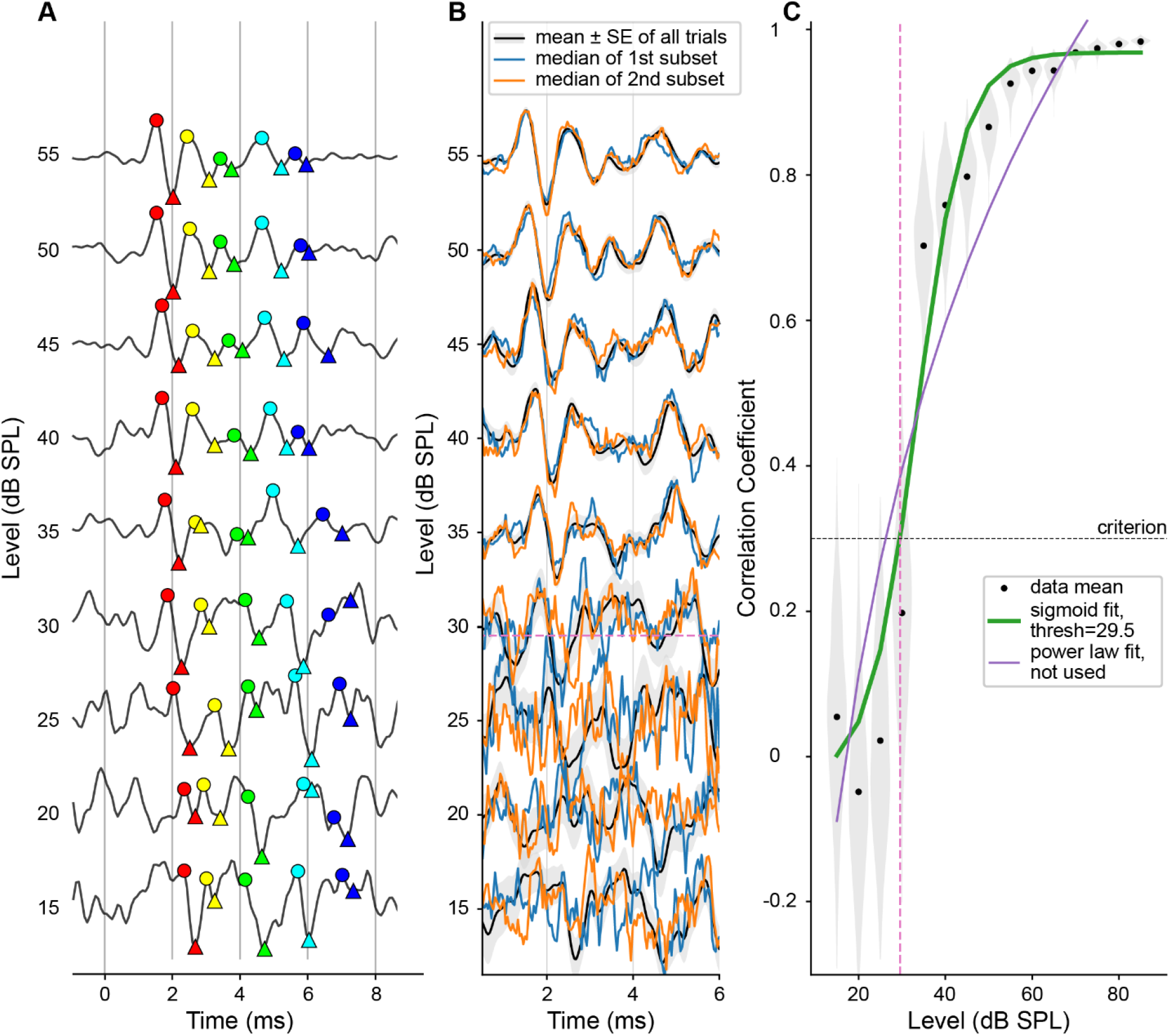
An example case of discrepancy between manual and algorithmic threshold. **A:** Stacked averaged ABR waveforms in response to 16-kHz tone-pips at levels surrounding threshold. Colored symbols represent peaks (circles) and troughs (triangles) of waves 1 to 5 identified by a human rater. **B**. Stacked ABR with mean ±SE and median of the two subsets (same representation is Figure 3A). **C**: Correlation coefficient as a function of level and curve fits (same representation as in Figure 3B).

There are a few other potential causes of “outliers” with larger discrepancies. In some cases, ABRpresto may overestimate the threshold due to noise obscuring the evoked response at near-threshold levels. On the other hand, in some cases ABRpresto reports thresholds below manual thresholds because ABR peaks are detected consistently across resamples. Because ABRpresto uses the median, it is more robust to large artifacts than human raters.

In order to find the threshold, the ABRpresto algorithm fits curves to determine when the normalized cross-correlation between subaverages is above a criterion level. Figure 6 shows the performance of the algorithm as a function of this criterion level. The percentage of algorithmic threshold judgements within ±5 dB and ±10 dB of manual judgements peaked at 69% and 92%, respectively. The performance in terms of the Spearman’s correlation coefficient was relatively flat at 0.97 for all values tested. Due to the less than optimal electrical conditions where background electrical noise levels can complicate the ABR interpretation, there can be variability in human judgements.

**Figure 6.**
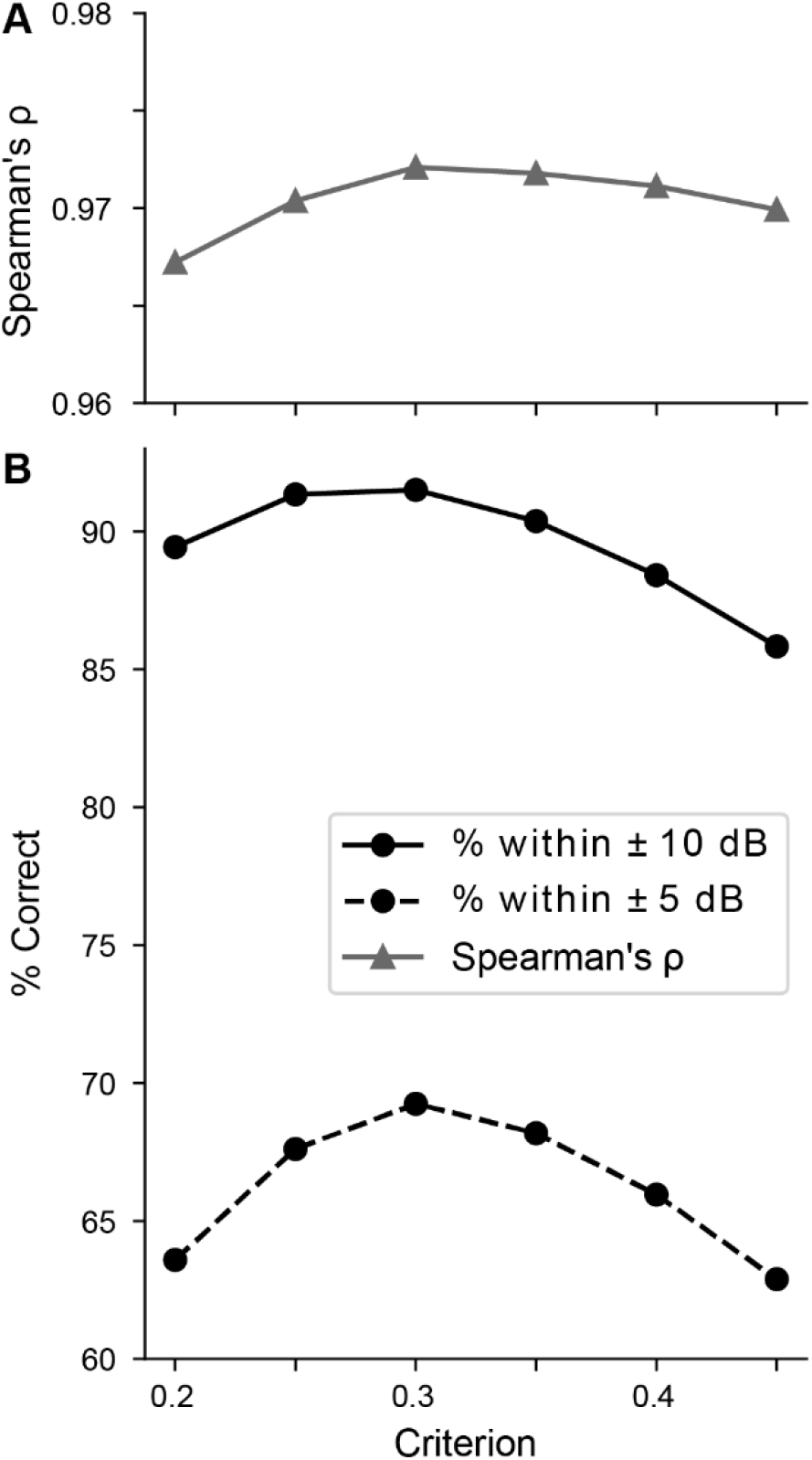
Sensitivity to criterion. **A**: Spearman’s ρ. Note tight axis limits, indicating the correlation coefficient is near 0.97 for all criteria. **B**: Percent of thresholds within ±10 dB and ±5 dB as a function of criterion for correlation coefficient.

Therefore, we favor maximization of the percentage of judgements within ±10 dB and chose a criterion of 0.3. Since the performance is relatively insensitive to the criterion, it is likely not critical to optimize this value for data acquired with other systems.

Figure 7 shows an efficacy study evaluating two different constructs of gene therapy candidates injected into a strain of mutant mice. The orange line and shaded area shows median and 90^th^ percentile range of ABR thresholds from wild-type control mice (C57BL/6), while the blue line and shaded area shows median and 90^th^ percentile range from untreated mutant mice. Pink and green lines represent thresholds from individual mice treated with two different gene therapy candidates. Both manual thresholds (Figure 7A) and algorithmic thresholds (Figure 7B) show that gene therapy 1 resulted in restoration of cochlear function to near normal level whereas gene therapy 2 produced partial restoration. The overall distribution of the thresholds based on manual vs. algorithmic thresholds is similar, leading to the same conclusion. However, the threshold curves of each individual mouse are smoother when using algorithmic thresholds. This is largely because of the algorithm’s ability to interpolate between tested levels, but also likely due to over- or under-estimation of human-rated manual thresholds due to noise. Therefore, the ABRpresto algorithm may increase the ability to infer differences between groups when the effect size is small.

**Figure 7.**
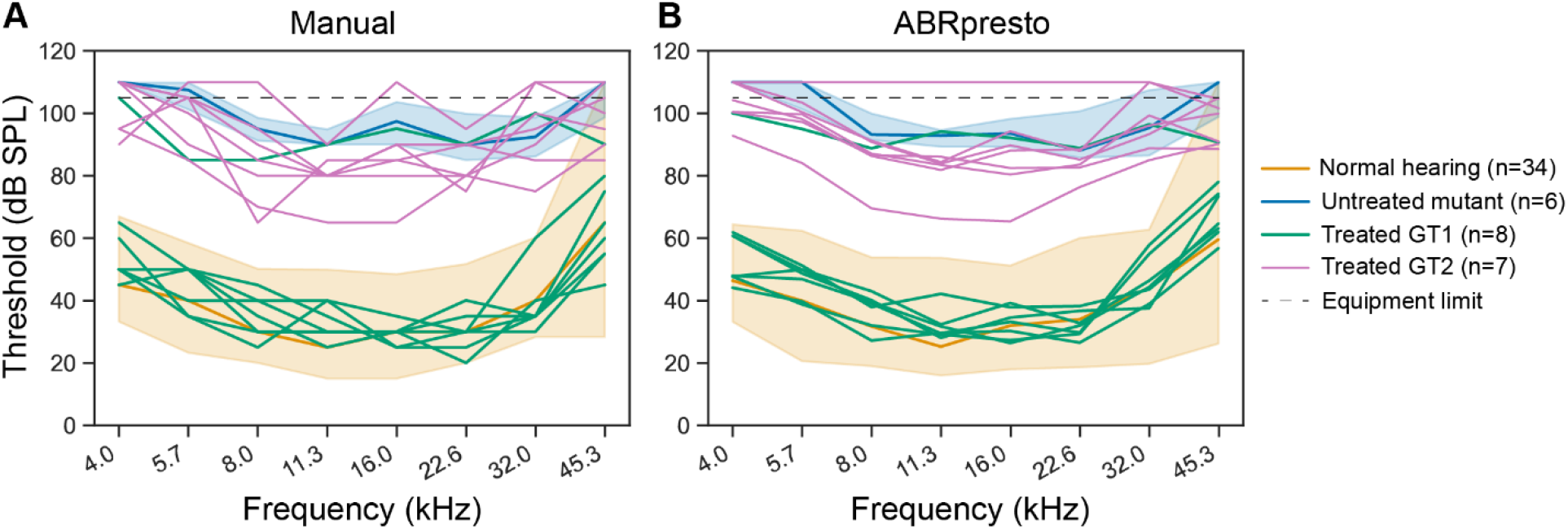
Manual vs algorithmic thresholds from a study evaluating two different gene therapy (GT) candidates in a mutant mouse model. ABR thresholds for normal hearing and untreated mutants are represented at median and 90th percentile range. **A**: Manual thresholds. **B**: Algorithmic thresholds.

## 4. Discussion

We developed a new algorithm for automatic estimation of ABR thresholds using resampled cross-correlation of two independent averages of responses to the same stimulus. The newly developed algorithm demonstrates robust and reliable performance, achieving 92% accuracy within ±10 dB of human-rated thresholds.

It is difficult to compare the performance of various existing algorithms because different metrics are used for evaluation. Here, we compare the current algorithm to select algorithms and directly compare the accuracy of two other algorithms based on cross-correlation on our data set (Table 1). The algorithm by Suthakar and Liberman (2019) employs cross-correlation of the average waveforms across levels. We re-analyzed this dataset and found the accuracy within ±10 dB to be 78% and 73% for the two observers of their dataset 1 (their Fig 6C and 6D) and 87% and 71% for the observers of their dataset 2 (their Fig 6E and 6F). We also applied this algorithm to the current dataset (of 7,857 thresholds) and found an accuracy of 78% of the reported thresholds within ±10 dB of human rated thresholds.

**Table 1.** ABRpresto performance compared to other algorithms. *Re-analysis of Suthakar and Liberman (2019) data, averaged result between two datasets and two human raters.

| Algorithm | Citation | Uses single-trial data? | Needs retraining for new datasets? | Reported in paper |  | Accuracy (on Decibel data) |
| --- | --- | --- | --- | --- | --- | --- |
|  |  |  |  | Large dataset? | Accuracy |  |
| Cross-correlation across levels | Suthakar and Liberman (2019) | No | Need to optimize criterion | Yes | Spearman's $\rho=0.8$<br>~77% within 10 dB* | 72% within 10 dB<br>Spearman's $\rho=0.95$ |
| Cross-correlation across subaverages within a level, time of peak | Wang et al. (2021) | No | Unknown | No, N=10 | Adjusted $R^2=0.99$ | 64% within 10 dB |
| Convolutional Neural Network | Thalmeier et al. (2022) | No | Yes | Yes | ~93% within 10 dB | Untested |
| "Sound Level Regression"<br>Smoothed low-frequency signal power | Thalmeier et al. (2022) | No | No | Yes | ~86% within 10 dB | Untested |
| ABRA: Convolutional Neural Network | Erra et al. (2024) | No | Possibly | Yes | 91% within 10 dB | Untested |
| **Resampled cross-correlation across subaverages within a level, mean amplitude** | **ABRpresto (this algorithm)** | **Yes** | **Need to optimize criterion** | **Yes** | **92% within 10 dB**<br>**Spearman's $\rho=0.97$**<br>**(reported in this paper, see Fig 4)** |  |

Like ABRpresto, the algorithm described by Wang et al. (2021) uses cross-correlation across subaverages. Rather than basing the threshold on the value of the cross-correlation at a lag of 0 ms, they define the threshold as the lowest stimulus level for which the cross-correlation functions have a peak within ∼1 ms of 0 lag. In a dataset of 10 threshold judgements, they reported an adjusted R^2^ of 0.99 and did not report percentage within ±10 dB. We re-implemented this algorithm and found an accuracy of 64% within ±10 dB on the current dataset. Thus, ABRpresto estimates thresholds more accurately than the two other algorithms based on cross-correlation.

Thalmeier et al. (2022) introduced two algorithms: one using a Convolutional Neural Network (CNN) and another based on sound level regression. The CNN has been reported to achieve approximately 93% accuracy within ±10 dB. The sound level regression algorithm also does not use single-trial data, does not need retraining, and achieves around 86% accuracy within ±10 dB. Erra et al. (2024) introduced 3 algorithms: a logistic regression, XGBoost, and a CNN. The CNN was the best performer, achieving 91% accuracy within ±10 dB. We did not evaluate the performance of these algorithms on our dataset. These algorithms have the advantage of not relying on single trial data, however, retraining may be needed for new datasets with highly diverse ABR waveform morphologies.

Given the reliance on human judgements to evaluate thresholding algorithms, the ABRpresto algorithm is likely close to optimal. Further refinement of curve fitting and edge cases may improve the performance slightly. Future work on algorithmic ABR thresholding could seek alternate strategies to identify a better “ground truth” upon which to compare top candidate algorithms and optimize. One possibility is to oversample threshold measurements by measuring responses to 10 times the normal number of repetitions in order to get a very clean signal for expert human judgement of thresholds, then compare that judgement to algorithm performance on a dataset with a normal number of repetitions. Another fruitful avenue may be to generate clean ABR waveforms with known thresholds using an auditory model and then computationally add noise and evaluate model performance. For both approaches, the challenge will be in maintaining representation of the vast array of waveform morphologies represented in various cases of pathology.

One challenge to the adoption of the ABRpresto algorithm is its use of single-trial data. Commercial ABR acquisition systems typically store only the averaged data (but in some cases there is an option to save single-trial data). We captured single-trial data using Psiexperiment, which also allowed us to use a stimulus repetition rate of 81 Hz using the interleaved ramp paradigm, allowing for rapid data collection (Buran et al., 2020). In some software that does not store single trials, separate averages in response to positive and negative polarity stimuli are stored. Unfortunately, these are likely not sufficient for the ABRpresto algorithm. In the current dataset, at least 10 resamples were needed to obtain a reasonably noise-insensitive estimate of the threshold. For datasets that do not contain single trials, the Suthakar and Liberman (2019), Thalmeier et al. (2022), or Erra et al. (2024) algorithms would all work well.

## 5. Conclusion

The newly developed algorithm has demonstrated good performance, achieving 92% accuracy within ±10 dB of human-rated thresholds. This validation was conducted using a comprehensive database comprising 7,857 thresholds across multiple strains and genotypes of mice with a wide range of hearing functions. Although it is challenging to define the “ground truth”, the new algorithm consistently produced the same conclusions as those derived from human ratings in all tested studies. As a result, this algorithm has now fully replaced the manual estimation of ABR thresholds for our preclinical studies, thereby saving significant time and enhancing objectivity in the process.

## Abbreviations

ABR: auditory brainstem response
CNN: convolutional neural network
DPOAE: distortion product oto-acoustic emissions
SPL: sound pressure level

## Acknowledgements

We thank Kathleen Berkun, Qyunh-Anh Artinian, Madison Traystman, and Sawanee Joshi for assistance in data acquisition, and Tara Robidoux and Sabrina Pray for animal care.

## Author contributions

Luke Shaheen: Conceptualization, Methodology, Investigation, Data curation, Formal analysis, Visualization, Writing – original draft, Writing – review and editing

Brad Buran: Conceptualization, Methodology, Software, Writing – review and editing Kirupa Suthakar: Formal analysis, Writing – review and editing

Seth Koehler: Conceptualization, Resources, Supervision, Project administration, Writing – review and editing

Yoojin Chung: Conceptualization, Investigation, Resources, Supervision, Project administration, Writing – original draft, Writing – review and editing

